# G protein-biased LPAR1 agonism promotes prototypic antidepressant effects

**DOI:** 10.1101/2022.11.02.514841

**Authors:** Naoto Kajitani, Mami Okada-Tsuchioka, Asuka Inoue, Kanako Miyano, Takeshi Masuda, Shuken Boku, Kazuya Iwamoto, Sumio Ohtsuki, Yasuhito Uezono, Junken Aoki, Minoru Takebayashi

## Abstract

Prototypic antidepressants, such as tricyclic/tetracyclic antidepressants (TCAs), have multiple pharmacological properties and have been considered to be more effective than newer antidepressants, such as selective serotonin reuptake inhibitors (SSRIs), in treating severe depression. However, the molecular mechanisms underlying the high efficacy of TCAs have not been completely understood. Herein, we found that lysophosphatidic acid receptor 1 (LPAR1), a G protein-coupled receptor, mediates the antidepressant effects of amitriptyline, a typical TCA. Amitriptyline directly bound to LPAR1 and activated downstream G protein signaling without affecting β-arrestin signaling, which implied that amitriptyline could act as a G protein-biased agonist of LPAR1. This biased agonism is unique to TCAs and has not been observed in other antidepressants, such as SSRIs. Long-term infusion of mouse hippocampus with 1-oleoyl-2-O-methyl-glycerophosphothionate (OMPT), a potent G protein-biased LPAR1 agonist, induced behavior similar to that induced by antidepressants. In contrast, LPA, a non-biased agonist of LPAR1, induced anxious behavior, indicating that LPAR1 may regulate conflicting emotional behaviors because of the downstream signaling bias. Furthermore, RNA-seq analysis revealed that LPA and OMPT have opposite patterns of gene expression changes in hippocampus. Ingenuity pathway analysis indicated that chronic intrahippocampal administration of OMPT could activate LPAR1 downstream signaling (Rho and MAPK), whereas LPA suppressed LPAR1 signaling. The results reveal the unique antidepressant effects of TCAs and indicate the potential of G protein-biased agonists of LPAR1 as targets for novel antidepressants.

## Introduction

Antidepressants have been developed to increase monoamine selectivity, as evidenced by serotonin and noradrenaline reuptake inhibitors (SNRIs) and selective serotonin reuptake inhibitors (SSRIs). While SNRIs and SSRIs have reduced side effects and increased tolerability compared to prototypic antidepressants, such as tricyclic/tetracyclic antidepressants (TCAs), their therapeutic effects may be comparable or even lower than those of TCAs (1, 2). Since approximately 30% of depression is treatment-resistant, antidepressants with pharmacological actions other than increasing monoamine selectivity need to be developed. Tricyclic antidepressants have been reported to be more effective than SSRIs in treating severe depression (2); however, mechanisms underlying their therapeutic effects remain unclear. Previous reports have suggested that in addition to modulating monoaminergic neurotransmission, TCAs may act on other therapeutic targets. In this study, we hypothesized that one of the potential targets of TCAs is the lysophosphatidic acid receptor 1 (LPAR1), a G protein-coupled receptor (GPCR). LPAR1 is one of the six receptors (LPAR1–6) that are activated by lysophosphatidic acid (LPA), a bioactive phospholipid. The receptor is activated by TCAs (3, 4) and contributes to emotional behaviors (5). LPAR1-deficient mice exhibited abnormalities in hippocampal functions and showed a phenotype with depressive and anxious features (6, 7). However, such a depression-like phenotype was also observed when LPA was administered into rodent brains (8, 9), which seems to contradict our hypothesis.

LPA binding to LPAR1 not only stimulates G protein signaling but also promotes receptor phosphorylation by G protein-coupled receptor kinases and subsequent binding of β-arrestins, which in turn mediates endocytosis and receptor desensitization (10). Although GPCRs typically activate both G protein signaling and β-arrestin-mediated endocytosis, some GPCR ligands preferentially activate either of the signaling pathways, a phenomenon referred to as biased agonism (11). For example, fingolimod, a β-arrestin-biased agonist of sphingosine 1-phosphate receptor, is a functional antagonist, since it selectively induces desensitization via β-arrestin-mediated endocytosis (12). LPA may cause functional antagonism of LPAR1 via endocytosis (13), and LPAR1 ligands that induce β-arrestin-mediated endocytosis may block LPAR1 signaling, as in the LPAR1-deficient mice. Therefore, we aimed to investigate whether LPAR1 could be involved in mediating the anti-depressive behavioral effects of TCA and to characterize the downstream signaling pathways of LPAR1 activated by TCAs.

## Results

### LPAR1 mediated the behavioral effects of TCA amitriptyline

In this study, amitriptyline was used as a typical TCA. We first determined whether LPAR1 is required for the behavioral effects of amitriptyline using the forced swim test (FST). Chronic treatment with amitriptyline decreased immobility time, indicating an antidepressant effect, which could be blocked by Ki16425, an LPAR1-3 antagonist (Fig. 1A to C). Moreover, LPAR1 heterozygous mice and a corticosterone-induced mouse model of depression (14) were used. Compared to null mice (15), LPAR1 heterozygous mice did not show neonatal mortality, weight loss, or abnormal behavior at baseline (Fig. S1). Similar to the results from Ki16425 treatment, LPAR1 heterozygous mice blocked the effects of chronic amitriptyline in the FST (Fig. 1D to F). Chronic administration of amitriptyline normalized the corticosterone-induced decrease in sucrose preference, an indicator of anhedonia, but had no effect in mice co-treated with Ki16425 (Fig. 1G to H). The results indicated that LPAR1 is involved in the behavioral changes induced by chronic amitriptyline treatment.

**Figure 1.**
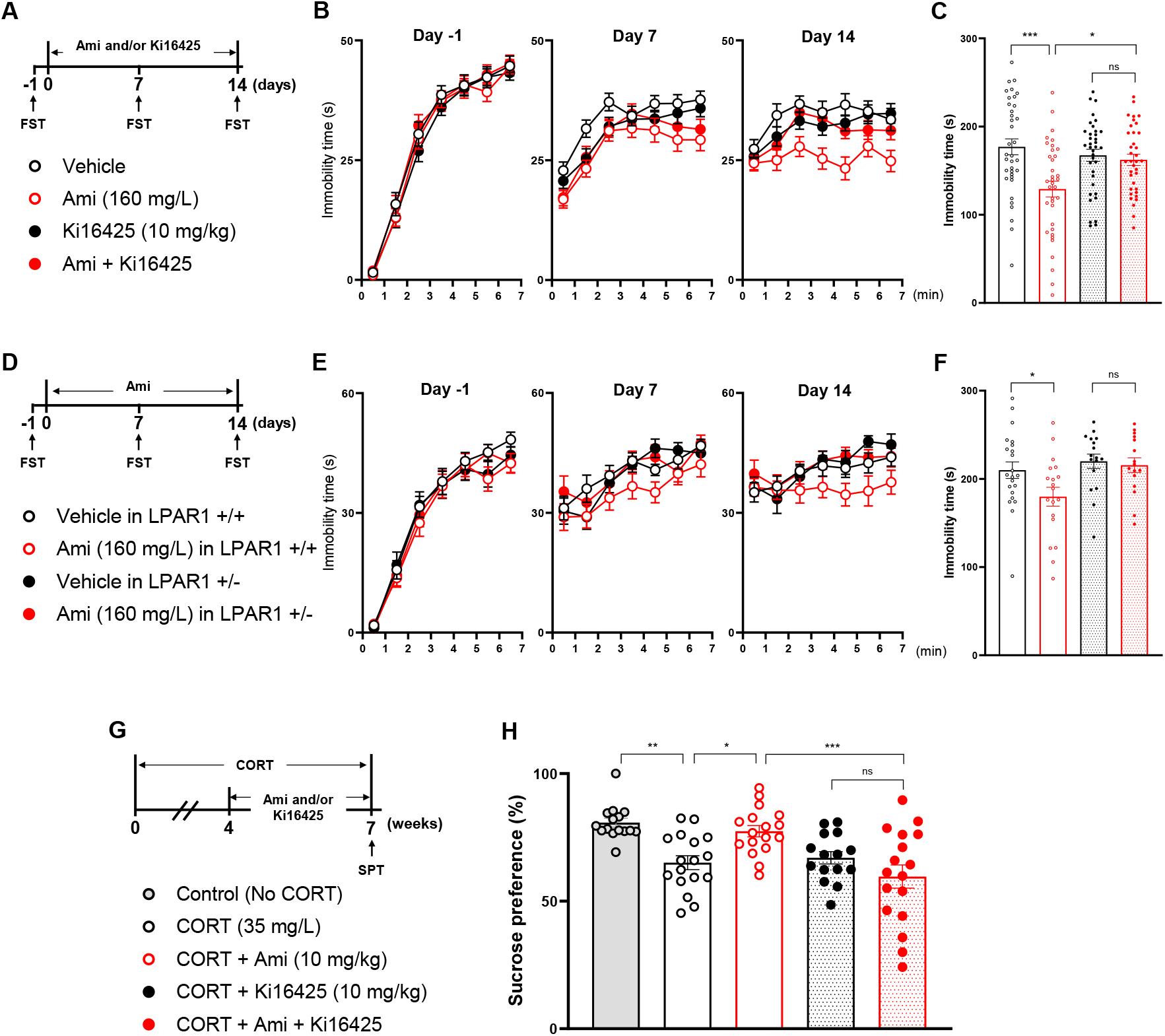
LPAR1 mediates the behavioral effects of amitriptyline (Ami). (A) Timeline and group design of the experiment are presented in panels B to C. (B) Immobility time at each minute in FST during three sessions (Day –1, 7, and 14) and (C) total immobility time during the last 5 min in FST on Day 14 in Ki16425 co-treated mice. (D) Timeline and group design of the experiment are presented in panels E to F. (E) Immobility time at each minute in FST during three sessions and (F) total immobility time during the last 5 min on Day 14 in LPAR1 heterozygous (+/-) mice. (G) Timeline and group design of the corticosterone (CORT)-treated mice. (H) Percentage of sucrose preference. Data are presented as means ± SEM. *P < 0.05, **P < 0.01, ***P < 0.001; see data S1 for complete statistics.

### TCAs directly bound to LPAR1 and competitively acted with LPA

We performed affinity purification using ferrite-glycidyl methacrylate beads (16) to determine whether TCAs directly bind to LPAR1. TCA-immobilized beads were synthesized by covalently attaching the beads to nortriptyline, a secondary amine derivative of amitriptyline (TCA-beads, Fig. 2A). Incubation of LPAR1-overexpressing cell lysate with TCA-beads eluted LPAR1 depending on the amount of immobilization (Fig. 2B to C). Free LPA or nortriptyline competitively inhibited the binding of LPAR1 to TCA-beads in a dose-dependent manner (Fig. 2D). Other free TCAs, such as amitriptyline or clomipramine, also inhibited the binding process (Fig. 2E), suggesting that LPA and TCAs bind directly to LPAR1.

**Figure 2.**
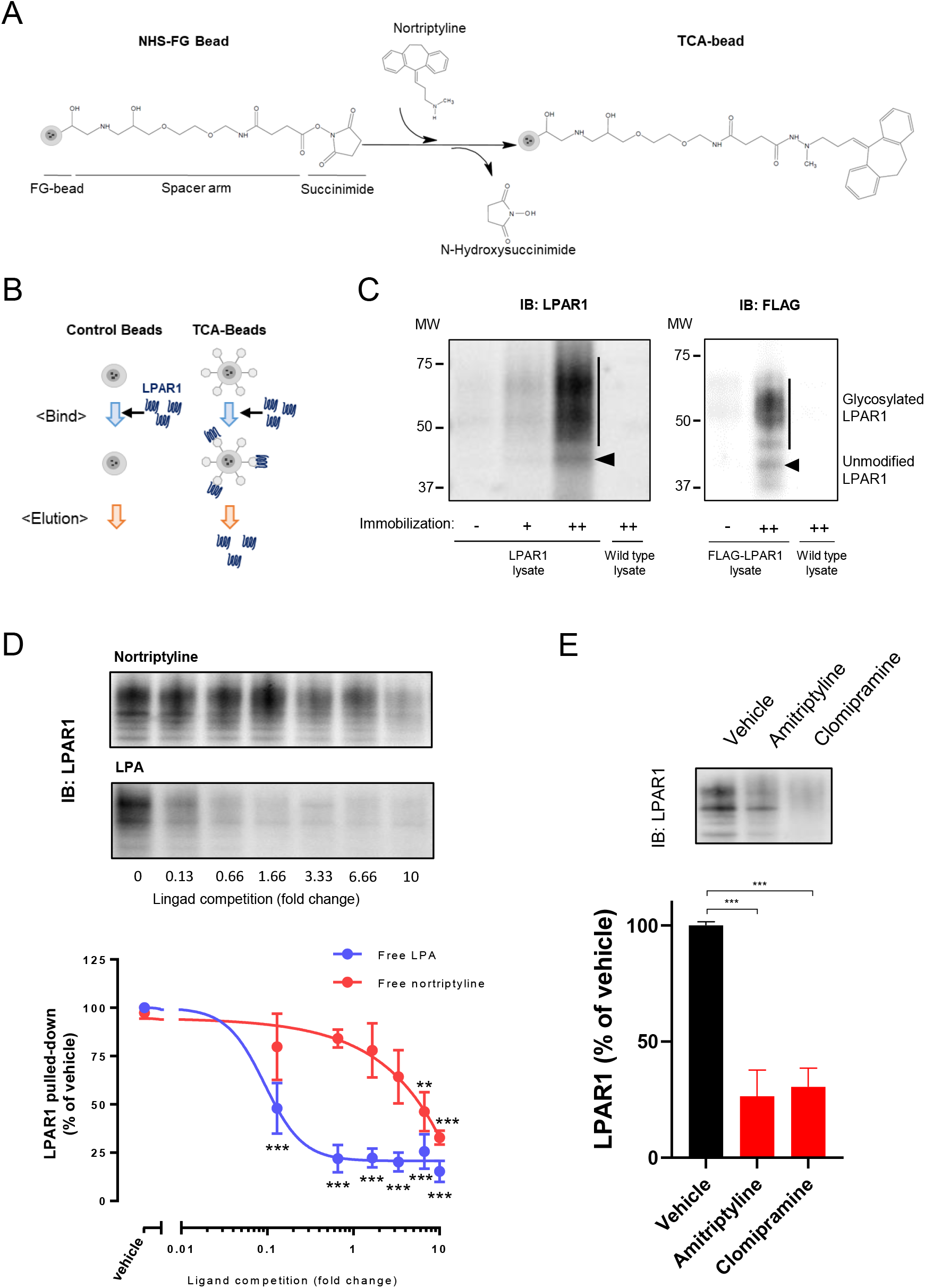
TCAs directly bind to LPAR1. (A) Scheme of nortriptyline immobilization on the FG beads. (B) Scheme of LPAR1 purification using TCA-beads. (C) Immunoblots from affinity-purified lysates using TCA-immobilized or control (-) beads. The amount of nortriptyline immobilized on the beads was 19.3 nmol/mg (+) or 27.3 nmol/mg (++). Two cell lysates were used for affinity purification, namely lysates from human LPAR1-overexpressed RH7777 cells (left) and those from FLAG-tagged human LPAR1-transfected HeLa cells (right). (D) Competitive inhibition of LPAR1 binding to TCA-beads. LPAR1-overexpressed RH7777 cell lysates were preincubated with the indicated concentrations of LPA or nortriptyline (shown as a ratio to the amount of nortriptyline immobilized on TCA-beads) and then eluted with TCA-beads. Representative immunoblots are shown above the graph. (E) Amitriptyline and clomipramine (6.67-fold concentration relative to the amount of nortriptyline immobilized on TCA-beads) inhibit LPAR1 binding to the TCA-beads. LPAR1-overexpressed RH7777 cell lysates were used. Representative immunoblots are shown above the graph. Data are presented as the means ± SEM of 3–4 independent experiments. *P < 0.05, **P < 0.01, ***P < 0.001; see data S1 for complete statistics.

A previous study using an electrical impedance (ΔZ)-based biosensor reported that amitriptyline activated G protein (increase in ΔZ) via endogenous LPAR1 in label-free cells (3). DAMGO, a ligand for µ-opioid receptors, which are GPCRs, also increased ΔZ in stable µ-opioid receptor-expressing HEK293 cells (Fig. S2A). Herein, amitriptyline additively increased the response of ΔZ by DAMGO, suggesting that amitriptyline and DAMGO are mediated by distinct receptors. In contrast, amitriptyline did not enhance the maximal response of ΔZ by LPA (Fig. S2B), hence indicating that amitriptyline and LPA act competitively on LPAR1.

### TCAs, but not other types of antidepressants, acted as a G protein-biased LPAR1 agonist

To investigate whether antidepressants have a signaling bias downstream of LPAR1, we measured G protein signaling and β-arrestin response via transforming growth factor α (TGFα) shedding assay (Fig. 3A) (17) and NanoBiT-based β-arrestin recruitment assay (Fig. 3B) (18), respectively. LPA was used as a reference. Amitriptyline treatment showed LPAR1-specific G protein signaling activity (E_max_ = 60.9 ± 4.1% and EC_50_ = 32.2 ± 8.5 µM), but less β-arrestin recruitment to LPAR1 (E_max_ = 8.8 ± 1.9%), indicating that amitriptyline is biased towards activating G protein signaling (Fig. 3C to E). Further, we examined whether different types of antidepressants could activate LPAR1 downstream pathways (Fig. S3). Considering that antidepressants can accumulate at tens of micromolar concentration in the brain (19, 20), antidepressants with E_max_ > 25% and EC_50_ < 100 µM were defined as agonists in this study. To quantitatively evaluate the potency of individual signaling activities of each agonist, we used relative intrinsic activity (RAi) values (21, 22). All tested TCAs showed agonist activity for G protein signaling, whereas the other tested antidepressant drugs, including SNRIs, SSRIs, ketamine, vortioxetine, or trazodone, did not (Fig. 3F). Mianserin and mirtazapine showed agonist activity for β-arrestin signaling but their G protein signaling activities were significantly higher or tended to be higher than their β-arrestin signaling activities (Fig. 3F). Results suggested that the G protein-biased agonism at LPAR1 is unique to TCAs.

**Figure 3.**
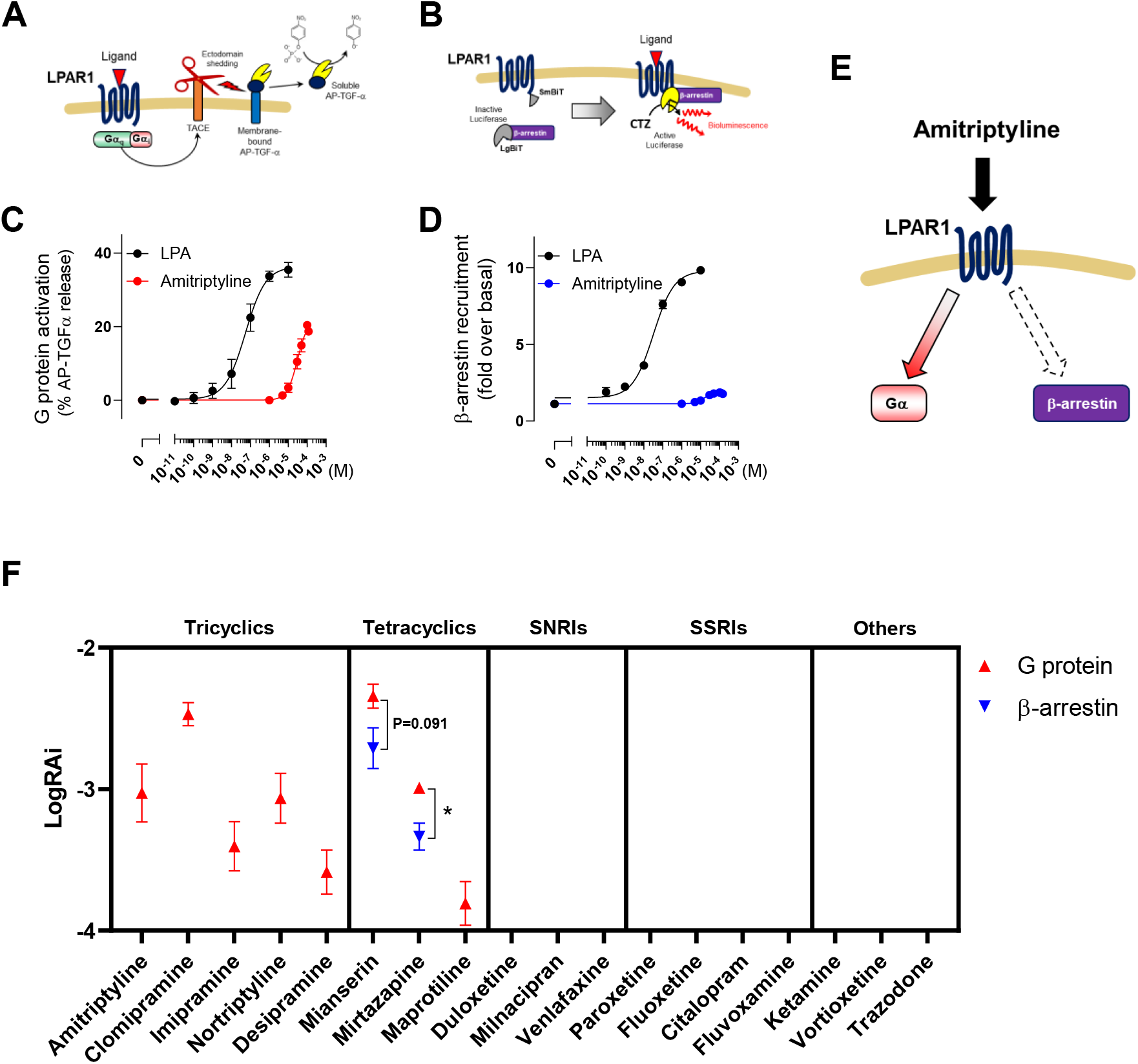
Tricyclic and tetracyclic antidepressants are G protein-biased LPAR1 agonists. (A) Schematic representation of TGFα shedding assay and (B) β-arrestin recruitment assay. Dose-response curves of LPA and amitriptyline for the signaling activities of (C) LPAR1-specific G protein and (D) β -arrestin. (E) Possible LPAR1 downstream pathways activated by amitriptyline. (F) LogRAi values for each antidepressant with agonist activity on the LPAR1-specific G protein and β -arrestin signal activities. Data are presented as means ± SEM of 3–4 independent experiments. *P < 0.05; see data S1 for complete statistics.

### Characterization of LPAR1 downstream signaling pathways using various LPAR1 agonists

We examined the effects of commercially available LPAR1 agonists, VPC31143, phospho-anandamide (pAEA), and 1-oleoyl-2-O-methyl-glycerophosphothionate (OMPT), on downstream signaling of LPAR1. They showed agonistic activities in both TGFα shedding and β-arrestin recruitment assays (Fig. 4A). Among the three agonists, OMPT showed a greater decrease in β-arrestin signaling than in G protein signaling (Fig. 4B). Immunoprecipitation confirmed that endogenous β-arrestin recruitment to FLAG-LPAR1 was not induced by OMPT (Fig. S4). The results overall suggested that OMPT is a potent G protein-biased LPAR1 agonist. LPA treatment decreased the cell surface expression of LPAR1 in HEK293 cells that lacked β-arrestin1/2 (Fig. S5A to B), hence suggesting that LPA induces LPAR1 endocytosis in a β-arrestin-dependent manner. In contrast, OMPT and amitriptyline did not induce β-arrestin-dependent endocytosis (Fig. S5C to D). Long-term treatment with LPA or OMPT did not decrease the protein level of FLAG-LPAR1 in HEK293 cells or LPAR1 protein in mouse hippocampus (Fig. S6). The results suggested that LPA reduces the cell surface expression of LPAR1 while retaining LPAR1 intracellularly without degrading it, consistent with previously reported findings (23), whereas OMPT does not reduce cell surface LPAR1 expression.

**Figure 4.**
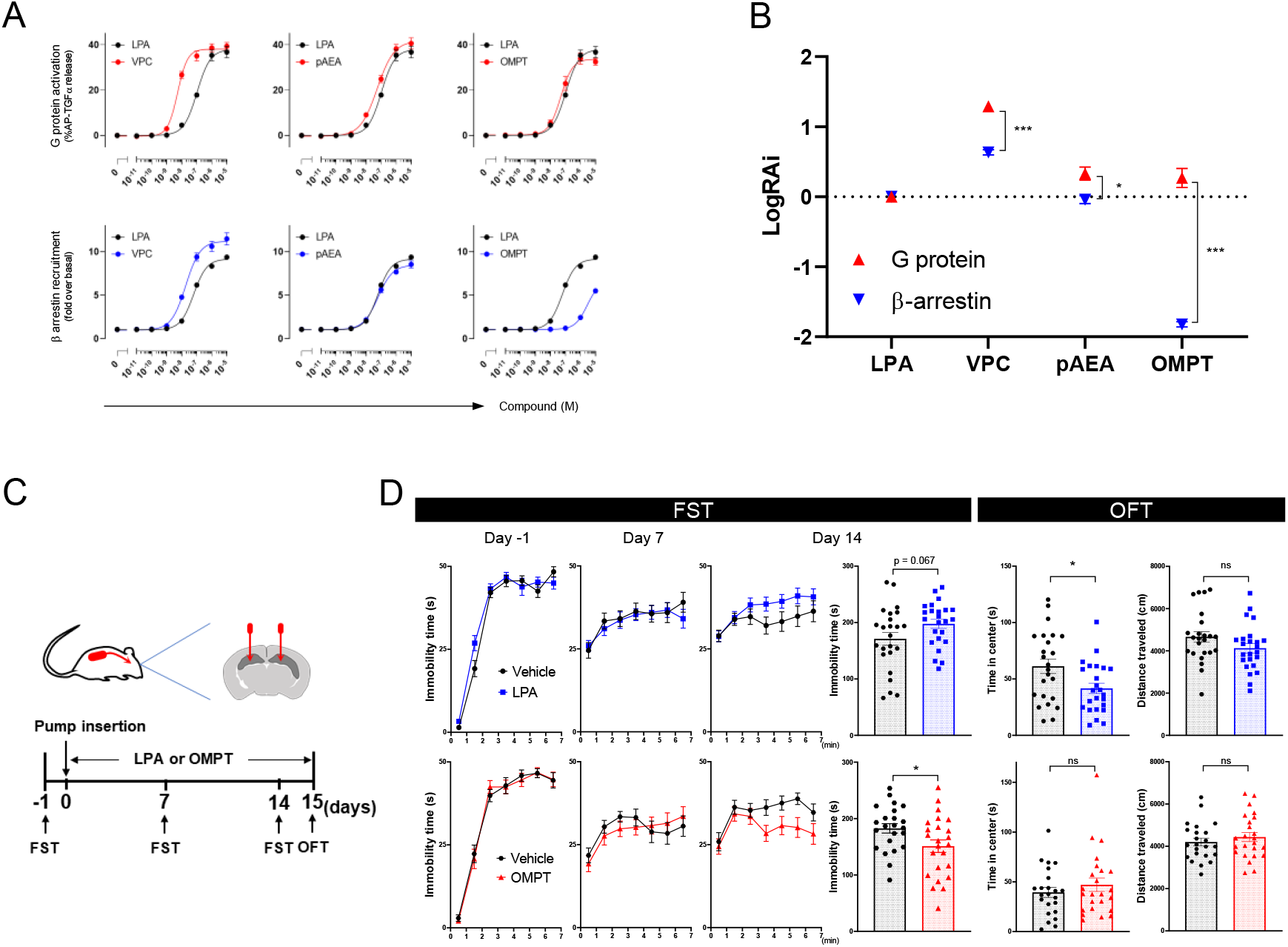
G protein-biased LPAR1 agonist induces antidepressant-like effects. (A) Dose-response curves and (B) logRAi values of LPAR1 agonists for LPAR1-specific G protein and β-arrestin signaling activities. (C) Timeline of the experiments using mice treated with intrahippocampal LPA or OMPT. (D) Immobility time at each min in FST during three sessions (Day –1, 7, and 14); total immobility time during the last 5 min in FST on Day 14, time in the center and distance traveled in the OFT. Data are presented as means ± SEM. *P < 0.05, ***P < 0.001; see data S1 for complete statistics.

### Chronic intrahippocampal infusion of OMPT, but not LPA, induced antidepressant-like effects

To investigate whether OMPT induces an antidepressant-like effect, OMPT was continuously infused directly into bilateral hippocampi using osmotic minipumps. Two weeks of OMPT infusion significantly decreased immobility time in the FST without affecting the locomotor activity in the open field test (OFT) (Fig. 4C to D). In contrast, consistent with previous reports (8, 9), infusion of LPA tended to increase immobility time in the FST, and decreased time spent at the center in the open field, indicating anxious behavior. The results suggested that LPA and OMPT may induce conflicting emotional behaviors owing to downstream signaling bias

### Characterization of LPAR2–6 downstream signaling pathways using various antidepressants and LPAR agonists

We examined LPAR2–6 activities and found that most, though not all, TCAs have G protein-biased LPAR2–3 agonism; however, there was no effect on LPAR4–6 activities (Fig. S7 to S11). OMPT was a potent G protein-biased agonist for all LPARs (Fig. S12). Therefore, the possibility that LPARs other than LPAR1 could be involved in antidepressant effects cannot be excluded. LPAR1 was remarkably expressed in mouse brain (Fig. S13), suggesting that LPAR1 is a major contributor to central LPA signaling.

### Long-term administration of LPA and OMPT showed different gene expression patterns

RNA-seq was performed to analyze gene expression changes in the hippocampus after long-term infusion with LPA or OMPT. Threshold free rank-rank hypergeometric overlap (RRHO) analysis (24, 25) was used to characterize the relationship between gene expression patterns by LPA and OMPT. RRHO revealed a substantial overlap of discordantly regulated genes between LPA and OMPT (Fig. 5A). Ingenuity pathway analysis (26) using the discordantly overlapping genes revealed four of the top five canonical pathways, predicted to be activated by OMPT, to be associated with downstream signals (Rho and MAPK) of LPAR1 (15) (Fig. 5B and S14). In contrast, the pathways predicted to be activated by LPA were negatively regulated by LPAR1 (27), suggesting that long-term infusion of OMPT may activate LPAR1 signaling, whereas LPA may suppress the same.

**Figure 5.**
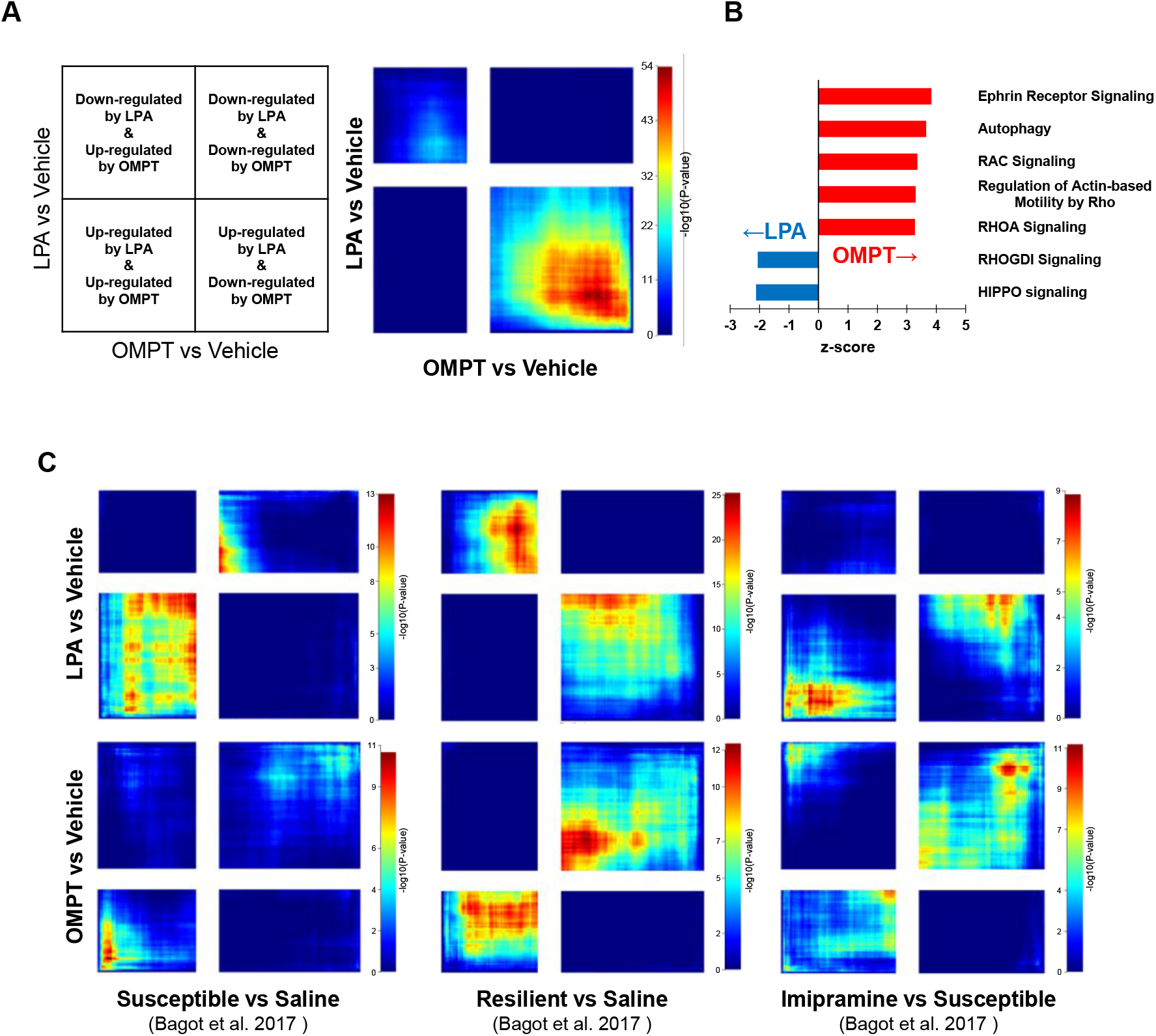
Transcriptional characterization by chronic administration of LPA or OMPT. (A) Comparison of RRHO expression patterns in the hippocampus between LPA and OMPT treatment. (B) Canonical pathways enriched with discordantly overlapping genes are presented in panel (A). Pathways with positive z-scores indicate pathways that are predicted to be activated by OMPT. (C) Comparisons of RRHO expression patterns in mice treated with LPAR1 agonists and in depression models treated with or without imipramine.

We used RRHO analysis to compare the present RNA-seq dataset with the published RNA-seq dataset of the chronic social defeat stress model and TCA-treated mice (28). Mice subjected to social defeat were divided into subpopulations that exhibited depression-like behaviors (susceptible) and those that did not (resilient). Depression-like behaviors in susceptible mice were ameliorated by the TCA imipramine (28). The transcription pattern of LPA-injected mice showed a concordant overlap with that of susceptible mice, but a discordant overlap with that of resilient mice (Fig. 5C). In contrast, the transcription pattern of OMPT-injected mice showed a concordant overlap with those of resilient and imipramine-treated mice, suggesting that gene expression patterns in OMPT-treated mice were similar to those in resilient and TCA-treated mice.

## Discussion

In the present study, we demonstrated that TCAs directly interact with LPAR1 and exhibit G protein-biased LPAR1 agonism, as implicated in the antidepressant effects. The agonist activity of LPAR1 was found to be unique to TCAs. Previous clinical findings have suggested that TCAs may be more effective than other types of antidepressants. A network meta-analysis of 21 antidepressants for acute phase treatment of major depressive disorder showed the TCA amitriptyline to be the most effective antidepressant (1). Another meta-analysis showed that tricyclic antidepressants are more effective than SSRIs in patients with severe depression (2). Tricyclic antidepressants confer higher rate of mood switching from depression to its opposite state of mania in patients with bipolar depression than other classes of antidepressants (29). The action of LPAR1 may partly explain such high responsiveness of TCAs. TCAs are less tolerated than SSRIs, owing to their several side effects, including antihistamine and anticholinergic effects. Thus, compounds with G protein-biased agonist activity of LPAR1 and monoamine neuronal activation could be new antidepressants with potential therapeutic effects.

We found an interesting phenomenon in which long-term treatment with LPAR1 agonists LPA and OMPT, which differ in their downstream signaling bias, induced conflicting emotional behaviors and regulated distinct gene expression patterns in hippocampus. LPA decreased the cell surface LPAR1 in a β-arrestin-dependent manner, suggesting that it may act as a functional antagonist. In contrast, G protein-biased LPAR1 agonists, OMPT and amitriptyline, did not induce β-arrestin-dependent endocytosis. Therefore, long-term administration of LPA might act as a functional antagonist and result in a behavior similar to that in LPAR1-deficient mice, whereas OMPT may be able to continue to activate LPAR1 and induce antidepressant-like effects. Thus, the concept of functional antagonism may provide a mechanism to resolve the previous inconsistent results in which LPAR1-deficient mice and LPA-treated mice both exhibited depression-like behaviors (6-9).

Recent structural studies indicated that GPCRs are highly dynamic proteins that can adopt multiple conformational states depending on their ligands (30, 31). Different receptor conformations result in different recruitment profiles for effector proteins, such as G proteins and β-arrestins. For example, G protein-biased agonists of µ-opioid receptor trigger conformational changes in the intracellular loop 1 and helix 8 domains of the µ-opioid receptor, thereby possibly impairing β-arrestin binding or signaling (32). The findings could allow for structure-based design of biased ligands. Recently, agonist-bound structures of human LPAR1 have been identified (33, 34). Therefore, future analysis of the conformational states of LPAR1 induced by OMPT and TCAs may yield important information for the discovery of novel antidepressant drugs.

Our findings implied that MAPK and Rho-related signaling, which are downstream signals of LPAR1, mediate antidepressant-like effects in the hippocampus. Previous studies had reported that neuronal Rho/ROCK inhibition causes antidepressant-like effects via neuronal morphological alterations (35, 36). Notably, LPAR1 has been reported to be heterogeneously expressed in the brain, with abundant expression in glial cells, such as oligodendrocytes and astrocytes, rather than in neurons (37). Since the Rho/ROCK pathway is important for glial cell proliferation (38), Rho signaling may play a distinct role in exhibiting antidepressant effects in LPAR1-expressing glial cells than in neurons.

In summary, our findings suggested that G protein-biased LPAR1 agonism may contribute to the non-monoaminergic antidepressant effects of TCAs. Further characterization of G protein-biased signaling of LPAR1 in the hippocampus can provide a basis for the development of novel antidepressants exhibiting activities other than monoamine restriction.

## Materials and Methods

### Animals

Male C57BL/6J mice and LPAR1 heterozygous mice (B6N(Cg)-Lpar1<tm1b(EUCOMM)Wtsi>/J, Stock No.27468) were used in this study; they were 7–8 weeks old at the beginning of the experiments. Mice were obtained from The Jackson Laboratory Japan (Yokohama, Japan) and maintained on a 12-h light/dark cycle with food and water ad libitum at a controlled temperature (23–25 °C). All experimental procedures were performed in accordance with the Guideline for Animal Experiments in National Hospital Organization Kure Medical Center and Chugoku Cancer Center (NHOKMCCCC). Protocols were approved by the Animal Research Ethics Committee, NHOKMCCCC.

### Behavioral procedures

The FST was performed in a clear acrylic cylinder (20 cm diameter), which was filled up to 16 cm with water (23–25 °C). Mice were placed in the cylinder and left there for 7 min. The entire session was videotaped, and immobility time was measured by the SMART video tracking system (SMART v3.0.06, Panlab, Barcelona, Spain).

The OFT was performed in a square chamber (36 × 36 × 30 cm) made of white polyvinyl chloride. Mice were placed around the novel open-field chamber and allowed to explore for 15 min. The entire session was videotaped. The time spent in the center of the chamber (18 × 18 cm) and the distance traveled were measured using the SMART video tracking system.

For the sucrose preference test (SPT), mice were exposed to drinking water substituted with 2% sucrose for three nights to avoid neophobia. Twenty-four hours before the SPT, mice were housed individually and subjected to water deprivation. Two pre-weighed bottles (one containing tap water and another containing 2% sucrose solution) were presented to each mouse for 4 h. The positions of water and sucrose bottles were switched 0.5 h and 2 h after the start of the SPT to avoid any place preference. The bottles were weighed again, and the difference in weight between 0.5–4 h was used to calculate the volume intake from each bottle. Sucrose preference was expressed as the percentage of sucrose intake relative to the total intake.

### Drug treatment timeline

For chronic amitriptyline treatment, amitriptyline was dissolved in tap water at a concentration of 160 mg/L. The average intake dose of amitriptyline per mouse during the experiment was 12.99 ± 0.19 mg/kg/day based on the average amount of water consumed and the average weight of the mice used in this study. Before amitriptyline administration, FST was performed to measure the basal immobility time. Mice were then grouped to ensure that there was no bias in basal immobility time. FST was performed at 1 and 2 weeks after administration.

For chronic corticosterone treatment, corticosterone (Tokyo Chemical Industry, Tokyo, Japan) was dissolved in 0.45% hydroxypropyl-β-cyclodextrin (β-CD; Fujifilm Wako, Osaka, Japan). Mice were treated with corticosterone (35 mg/L) or 0.45% β-CD, added to their drinking water, for 7 weeks. The average intake dose of corticosterone per mouse during the experiment was 7.29 ± 0.68 mg/kg/day. During the last 3 weeks of corticosterone treatment, mice were daily injected with amitriptyline (10 mg/kg/day, i.p.) and/or Ki16425 (10 mg/kg/day, i.p.), and SPT was performed subsequently.

For continuous intrahippocampal injection, mice were anesthetized with isoflurane and surgically implanted with two subcutaneous osmotic minipumps (Alzet model 1004; Durect Corporation, Cupertino, CA, USA) and bilateral guide cannulae (Plastics One, Roanoke, VA, USA) targeting hippocampi. The minipumps were filled with LPA (15 nM in PBS), OMPT (15 nM in 40% DMSO and 60% PBS), PBS (vehicle for LPA), or 40% DMSO-PBS (vehicle for OMPT), and activated the evening before surgery by incubating them at 37 °C in saline to initiate a continuous delivery at 0.11 µL/h over 2 weeks. Bilateral cannulae were delivered into the hippocampus at - 2.2 mm posterior to the bregma, ± 1.5 mm lateral to the midline, and - 2.0 mm ventral to the surface of the skull. The antibiotic penicillin G (500 units/mouse, i.m.) and the analgesic carprofen (5 mg/kg, i.p.) were administered after surgery. Before the surgery, FST was performed to measure basal immobility time. Mice were then grouped to ensure that there was no bias in basal immobility time. FST was performed at 1 and 2 weeks after the surgery, followed by OFT.

### Preparation of tricyclic antidepressant (TCA)-beads

Nortriptyline was fixed on NHS-beads (TAS8848 N1141; Tamagawa Seiki, Nagano, Japan) according to the manufacturer’s instructions. To prepare different fixed levels of TCA-beads, 0.3 mM or 0.5 mM nortriptyline was incubated with NHS-beads. Non-fixed beads were taken as control beads. The amount of nortriptyline immobilized on beads was calculated by quantifying the N-hydroxysuccinimide, cleaved by the ligand immobilization reaction (see Fig. S2A), using high-performance liquid chromatography (outsourced to Tamagawa Seiki). The average amount of fixation was 19.3 nmol/mg using 0.3 mM nortriptyline and 27.3 nmol/mg using 0.5 mM nortriptyline.

### Affinity purification of LPAR1 with TCA-beads

LPAR1-overexpressing membrane lysates derived from RH7777 were obtained from Chantest (#A324; Cleveland, OH, USA), and prepared from HeLa cells (obtained from RIKEN BRC, RBRC-RCB0007) transfected with FLAG-LPAR1 plasmid. HeLa cells were transfected with FLAG-tagged human LPAR1-encoding pCAGGS plasmid using ScreenFectA plus and the enhancer (SFA P-reagent; Fujifilm Wako), according to the manufacturer’s instructions. The lysates were incubated with control beads or TCA-beads for 2 h at 4 °C in binding buffer (20 mM HEPES-NaOH, pH 7.9, 100 mM NaCl, 10 mM MgCl_2_, 10% glycerol, 0.2% NP-40) with protease inhibitor cocktail (Sigma). Magnetic separation and washing were repeated four times with wash buffer (20 mM HEPES-NaOH, pH 7.9, 100 mM NaCl, 10 mM MgCl_2_, 10% glycerol, 0.1% NP-40). The binding proteins were eluted by adding 1× SDS sample buffer and boiled at 100 °C for 5 min. In competitive experiments, free ligands were pre-mixed with the lysate and incubated with the TCA-beads. Eluted LPAR1 was detected by immunoblotting, as described below.

### Immunoblotting

Immunoblotting was performed with individual antibodies, as follows: anti-EDG2 antibody (for LPAR1) (ab23698, Abcam, Cambridge, MA, USA), anti-DDDDK tag antibody (for FLAG) (PM020, MBL, Tokyo, Japan), anti-FLAG (M2) antibody (F1804, Sigma, St.Louis, MO, USA), anti-β-arrestin1/2 antibody (sc-74591, Santa Cruz, CA, USA), and anti-Na^+^/K^+^-ATPase antibody (#3010, Cell signaling, Danvers, MA, USA). Each antibody was diluted at an appropriate concentration with Can Get Signal Solution (Toyobo, Osaka, Japan) and treated with the blotting membrane.

Protein amount-adjusted lysate and eluted sample in bead affinity purification or immunoprecipitation were separated by SDS-polyacrylamide gel electrophoresis and transblotted onto polyvinylidene difluoride membranes. The membranes were blocked with 5% nonfat dry milk in TBST for 1 h at room temperature. Thereafter, the membranes were incubated with primary antibodies overnight at 4 °C. After washing, the membranes were incubated with horseradish peroxidase-conjugated secondary antibodies for 1 h at room temperature. Chemiluminescence detection was performed using Immun-Star WesternC Kit (Bio-Rad, Hercules, CA, USA), and the net intensities of each signal were quantified using ChemiDoc Imaging Systems (Bio-Rad).

### Transforming growth factor α (TGFα) shedding assay

The TGFα shedding assay, which measures the activation of specific GPCR-dependent G protein signaling, was performed as described previously (17), with minor modifications. HEK293FT cells (obtained from Life Technologies, cat. no. R700-07) were seeded in 60-mm culture dishes (8 × 10^5^ cells/dish) in DMEM (Nissui Pharmaceutical, Tokyo, Japan) supplemented with 10% fetal bovine serum, glutamine, penicillin, and streptomycin (growth medium). After a 1-day culture, cells were transfected with the transfection solution that was prepared by combining 8 µL (per dish, hereafter) polyethyleneimine (PEI) solution (1 mg/mL; Polysciences, Warrington, PA, USA) and a plasmid mixture consisting of 400 ng human LPAR-encoding plasmids, 1000 ng alkaline phosphatase (AP)-tagged TGFα (AP-TGFα)-encoding plasmid with or without 200 ng chimeric Gα subunit protein (Gαq/i1 for LPAR1, LPAR5, and LPAR6, and Gαq/s for LPAR2 and LPAR4)-encoding plasmid in 400 µL of Opti-MEM (Thermo Fisher Scientific, Waltham, MA, USA). After a 1-day culture, the transfected cells were harvested by trypsinization, neutralized with the growth medium, and collected by centrifugation at room temperature at 200 g for 5 min. Cells were suspended in HBSS containing 5 mM HEPES (pH 7.4) and were left for 10 min to remove extra AP-TGFα released during trypsinization. After centrifugation, cells were resuspended in 12 mL of HEPES-HBSS and seeded in a 96-well plate (90 µL/well). Cells were incubated at 37 °C for 30 min to allow the cells to attach. Test compounds were diluted in 0.01% fatty-acid-free grade BSA (Fujifilm Wako)-containing HEPES-HBSS (assay buffer) and added to the cells (10 µL/well). For LPAR4–6, 10 µM Ki16425, an LPAR1–3 antagonist, was pretreated 5 min before the addition of test compounds. After a 1-h incubation, cells in 96-well plates were centrifuged, and the conditioned medium (80 µL) transferred to an empty 96-well plate. The AP reaction solution (a mixture of 10 mM p-nitrophenyl phosphate, 120 mM Tris-HCl (pH 9.5), 40 mM NaCl, and 10 mM MgCl_2_) was added to plates containing cells and the conditioned medium (80 µL/well). Absorbance at a wavelength of 405 nm was measured using a microplate reader (Varioskan LUX multimode microplate reader, Thermo Fisher Scientific) before and after 1-or 2-h incubation at room temperature. All values were calculated by subtracting the background activity induced by compounds in empty plasmid-expressing HEK293FT cells. The AP-TGFα release percentages were fitted to a four-parameter sigmoidal concentration-response curve, using the GraphPad Prism 8 software (v8.4.3, GraphPad Software, San Diego, CA, USA), and the EC_50_ and E_max_ values were obtained therefrom. The E_max_/EC_50_ values of each compound were normalized by the E_max_/EC_50_ value of LPA to calculate RAi, which was then base-10 log-transformed (LogRAi) and used as the potency of signaling activation.

### NanoBiT-based β-arrestin1 recruitment assay

NanoBiT PPI assay-based β-arrestin1 recruitment assay was performed as described previously (39), with minor modifications. Human full-length β-arrestin1 was N-terminally fused to a large fragment (LgBiT; forming LgBiT-β arr1) of NanoBiT luciferase with a 15-amino-acid flexible linker (GGSGGGGSGGSSSGG). Human LPAR1–6 were C-terminally fused to a small fragment (SmBiT; forming LPARs-SmBiT) with the 15-amino-acid flexible linker. The LgBiT-β arr1 and the LPARs-SmBiT constructs were inserted into a pCAGGS expression plasmid vector. Transfection into HEK 293FT cells was performed as described in the TGFα shedding assay using the PEI method (200 ng LgBiT-β arr1, 1000 ng LPAR-SmBiT, and 8 µL of 1 mg/mL PEI solution per dish). After a 1-day culture, transfected cells were collected with 0.5 mM EDTA-containing PBS, centrifuged, and suspended in 2 mL of assay buffer. The cell suspension was seeded in a white 96-well plate at a concentration of 5 × 10^5^ cells/mL (80 µL/well) and loaded with 20 µL of 50 µM coelenterazine (Cayman, Ann Arbor, MI, USA) diluted in the assay buffer. After 2-h incubation at room temperature, the plate was measured (0.5 s/well) for baseline luminescence (Varioskan LUX multimode microplate reader) and 20 µL of test compounds diluted in the assay buffer were manually added. The plate was read 30 times (0.18 s/well) at room temperature. The luminescent signal was normalized to the initial count, and fold-change values over 5–10 min after test compound stimulation were averaged. The fold-change β -arrestin recruitment signals were fitted to a four-parameter sigmoidal concentration-response, and the EC_50_, E_max_, and LogRAi values were obtained as described above.

### RNA-seq and data analysis

To limit the contribution of acute effects of behavioral testing on gene expression, mice were sacrificed 2 days following the final behavioral test. Whole hippocampi were rapidly isolated and frozen in liquid nitrogen. Total RNA was extracted using the AllPrep DNA/RNA mini kit (Qiagen, Hilden, Germany) with DNase I to eliminate any contaminating DNA. The RNA integrity number values of all samples were above 8.0. Library preparation and RNA-seq were outsourced to Macrogen Japan (Kyoto, Japan). Libraries were prepared using the TruSeq Stranded mRNA library prep kit (Illumina, San Diego, CA, USA), and were sequenced on the NovaSeq 6000 (Illumina) with 100 bp paired-end reads.

Raw reads were trimmed for adaptor sequences and low-quality reads using Trimmomatic (v0.39) with the following parameters: LEADING, 20; TRAILING, 20; CROP, 100; MINLEN, 20. Trimmed reads were then aligned to the mm10 reference genome using STAR (v2.7.9a) with default parameters. Uniquely mapped reads were used to obtain the estimated read count and transcripts per million (TPM) using RSEM (v1.3.3). Estimated read count was processed using TCC-GUI (40) with default parameters. TCC-GUI uses the TMM normalization method, and edgeR was used to determine the differentially expressed genes. Before processing with TCC-GUI, poorly expressed genes were filtered by removing those with an average TPM value of 0 and those with an average TPM value in the bottom 25% of the remaining genes.

To evaluate the overlap of expression patterns between two ranked gene lists, threshold-free genome-wide transcriptomic overlap analysis was conducted using rank-rank hypergeometric overlap (RRHO2, v1.0). Transcripts from each TCC-GUI comparison were ranked by -log10 (p-value) multiplied by the sign of fold change. The ranked gene lists were used to generate matrices of overlapped genes between differential expression lists of interest. Discordantly overlapped genes between LPA and OMPT-elicited transcriptional patterns were extracted from the data analyzed by RRHO2. Using the gene lists, canonical pathways were generated through the use of Ingenuity Pathway Analysis (IPA, Release Date: 2021-10-22, Qiagen).

A publicly available RNA-seq dataset (Gene Expression Omnibus database: GSE81672) was obtained for comparison with our RNA-seq data. The data were processed using the pipeline described above and analyzed by RRHO2.

### Statistical analysis

All data are presented as mean ± SEM. Statistical significance was determined using the unpaired t-test, one-way ANOVA with post hoc Tukey’s multiple comparison test or Dunnett’s multiple comparisons test, two-way ANOVA with post hoc Tukey’s multiple comparison test, or mixed-effects model with post hoc Sidak’s multiple comparisons test using GraphPad Prism 8 software. In the figures, significant effects are denoted by asterisks that indicate *P < 0.05, **P < 0.01, and ***P < 0.001. Complete statistical information can be found in Data S1.

## Acknowledgments

We thank K. Hashimoto (Chiba University) for providing (R, S)-ketamine; N. Maeda and K. Matsuura (Kumamoto University) for assisting with experiments; Y. Nakachi (Kumamoto University) for helpful comments on this manuscript; and S Khanna from the Editage group for editing a draft of this manuscript.

